# Harmonic brain modes: a unifying framework for linking space and time in brain dynamics

**DOI:** 10.1101/162040

**Authors:** Selen Atasoy, Gustavo Deco, Morten L. Kringelbach, Joel Pearson

## Abstract

A fundamental characteristic of spontaneous brain activity is coherent oscillations covering a wide range of frequencies. Interestingly, these temporal oscillations are highly correlated among spatially distributed cortical areas forming structured correlation patterns known as the resting state networks, although the brain is never truly at ‘rest’. Here, we introduce the concept of **“harmonic brain modes”** – fundamental building blocks of complex spatiotemporal patterns of neural activity. We define these elementary harmonic brain modes as harmonic modes of structural connectivity; i.e. connectome harmonics, yielding fully synchronous neural activity patterns with different frequency oscillations emerging on and constrained by the particular structure of the brain. Hence, this particular definition implicitly links the hitherto poorly understood dimensions of space and time in brain dynamics and its underlying anatomy. Further we show how harmonic brain modes can explain the relationship between neurophysiological, temporal and network-level changes in the brain across different mental states; (*wakefulness, sleep, anaesthesia, psychedelic*). Notably, when decoded as activation of connectome harmonics, spatial and temporal characteristics of neural activity naturally emerge from the interplay between excitation and inhibition and this critical relation fits the spatial, temporal and neurophysiological changes associated with different mental states. Thus, the introduced framework of harmonic brain modes not only establishes a relation between the spatial structure of correlation patterns and temporal oscillations (linking space and time in brain dynamics), but also enables a new dimension of tools for understanding fundamental principles underlying brain dynamics in different states of consciousness.

## Introduction

Science aims to identify certain laws, rules or fundamental principles, which are consistently observed among interacting elements in natural phenomena, and to express them with mathematical equations. When it comes to understanding the brain, however, neuroscience is yet to discover a fundamental principle linking the brain’s *structure*, *function* and *subjective experience*.

So far, the most studied fundamental characteristic of mammalian brain activity is coherent oscillations covering a wide range of frequency bands ranging from 0.05 Hz to 500 Hz. (Buzsaki, 2004). Furthermore, it has also been long known that various states of consciousness, such as awake state, sleep and anesthesia are accompanied by changes in the frequency and amplitude of these oscillations; e.g. the low amplitude high frequency alpha oscillations mainly observed in conscious awake state turn into low frequency high amplitude delta waves in deep sleep stages (Steriade, McCormick, & Sejnowski, 1993). However, recent studies indicate that the relation between states of consciousness and changes in neural oscillations are more complex than a simple frequency-state relation (Massimini, 2005; Purdon et al., 2013). Crucially, changes in the state of consciousness, induced by sleep (Boly et al., 2012; Buckner, Krienen, & Yeo, 2013; Massimini, 2005; Tononi & Koch, 2008), anaesthesia (Alkire, Hudetz, & Tononi, 2008; Boly et al., 2012; Boveroux et al., 2010; Deco, Jirsa, & McIntosh, 2011; 2013b; Deco, Jirsa, McIntosh, Sporns, & Kötter, 2009b; Horovitz et al., 2009; Purdon et al., 2013; Tononi & Koch, 2008) or psychedelics (Carhart-Harris, Erritzoe, et al., 2012a; Tagliazucchi et al., 2014; Tagliazucchi et al., 2016, Carhart-Harris, Muthukumaraswamy, et al., 2016b), are accompanied by rather widespread changes in the coherent oscillations throughout the cortex (Deco, Hagmann, Hudetz, & Tononi, 2013a; Jobst et al., 2017).

A decade ago, a remarkable discovery revealed that temporal oscillations in spontaneous activity during awake resting-state, in fact exhibit highly structured correlation patterns among spatially distributed cortical regions (Biswal, Yetkin, Haughton, & Hyde, 1995; Damoiseaux et al., 2006; Deco & Corbetta, 2011; Fox, Snyder, Vincent, Corbetta, & Van Essen, 2005). This discovery created a paradigm shift in the neuroscience community from using task-based paradigms to studying spontaneous neural activity in the resting-state (Fox & Raichle, 2007). Notably, the topography of the observed correlation patterns, termed resting state networks (RSNs) (Boly et al., 2012; Deco et al., 2011; Fox et al., 2005; Fox & Raichle, 2007; Stewart, 1999), closely resembles the functional networks of the human brain identified by various sensory, motor, and cognitive tasks (Barttfeld et al., 2015; Buckner et al., 2013; Chladni, 1830; Deco et al., 2011) (see Figure 1). Although the overlap of the RSNs with functional networks of the human brain indicate their relevance for active cognitive processes, correlated low-frequency fluctuations of BOLD signal have been observed under a wide range of physiological conditions, including awake state (Biswal et al., 1995; Fransson, 2005), light sleep (Horovitz et al., 2008; Larson-Prior et al., 2009), anesthetised humans (Greicius et al., 2008) and anesthetised non-human primates (Vincent et al., 2007), introducing an apparent paradox regarding RSNs relationship to conscious awareness.

**Figure 1.**
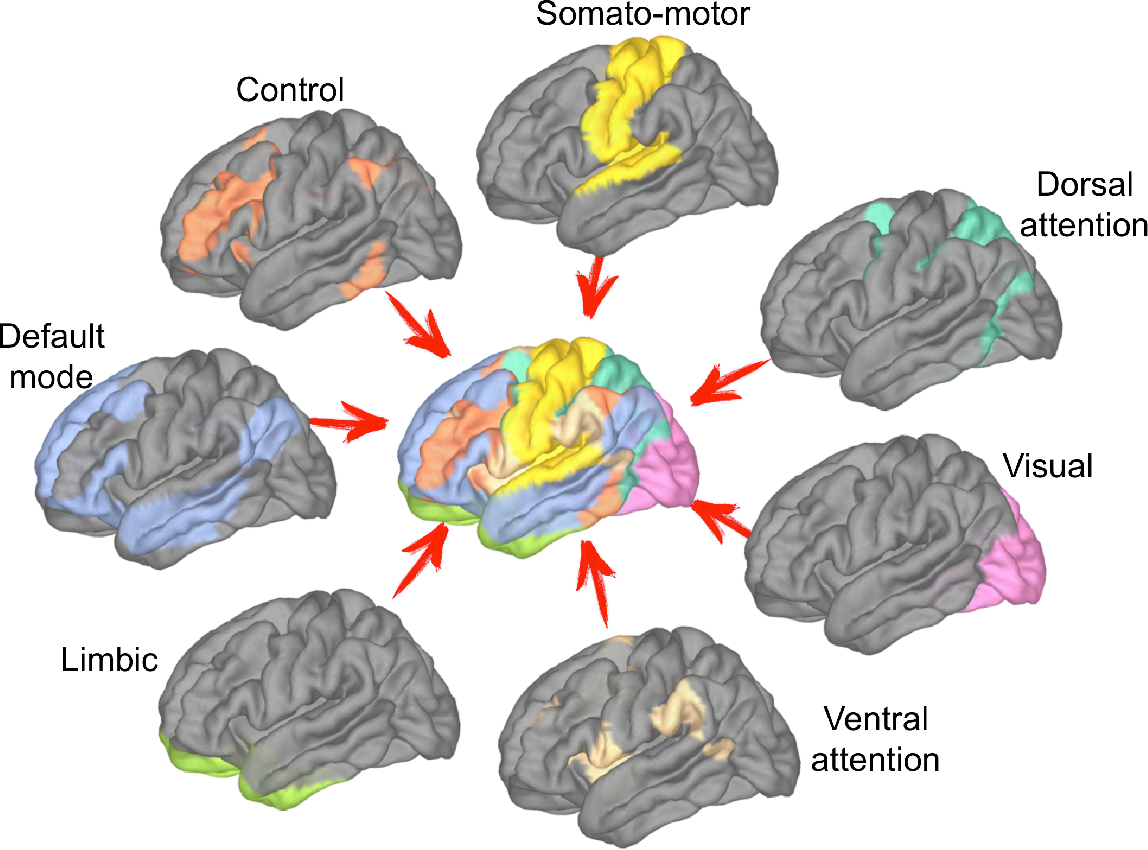
Resting state networks (RSNs) of the human brain overlap with the functional networks identified with various task-based cognitive paradigms. The image was reproduced using the networks published in (Thomas Yeo etal., 2011).

Crucially, a solution to this paradox and a full appreciation of RSNs’ role for brain function and subjective mental states requires an understanding of how they are linked to 1. *temporal oscillations* - a fundamental characteristic of mammalian brain activity (Buzsaki, 2004); 2. *human connectome* - the particular structural connectivity of the human brain (DeFelipe, 2010), and 3. *neurophysiology* - the interplay between neural excitation and inhibition. Although current developments in structural and functional neuroimaging, combined with computational models (Cabral, Kringelbach, & Deco, 2017; Deco, Jirsa, McIntosh, Sporns, & Kotter, 2009a; Honey et al., 2009) have shed light on the links between the RSNs, brain anatomy and neurophysiology, *
**integrating the dimensions of space and time in neural dynamics and discovering their relation to subjective mental states remains one of the biggest challenges in neuroscience**
*.

A critical starting point in this endeavour is to identify possible elementary brain modes – the macro-scale building blocks of neural activity - composing the complex spatiotemporal patterns that we measure in functional neuroimaging. Just like the different combinations of chemical basis in DNA comprise the fundamental elements of different genes or elementary particles form the building blocks of various atoms and molecules, here we introduce the concept of *harmonic brain modes*, as the fundamental building blocks of complex spatiotemporal neural activity patterns.

In this review article, we show how existing findings on a network-level, temporal and neurophysiological correlates of various *mental states (waking consciousness, sleep, anaesthesia, psychedelic states)* can be unified and understood within one simple mathematical framework that decodes brain activity as a combination of harmonic brain modes. We show how the harmonic brain modes can be (pre)-determined as the harmonic modes of the structural connectivity yielding fully synchronous neural activity patterns with different frequency oscillations (Atasoy, Donnelly, & Pearson, 2016). We describe the evidence in favour of the characterization of spontaneous activity in various mental states in terms of these harmonic brain modes. Furthermore, we relate the mathematical framework of these elementary brain modes to the fundamental principle of harmonic patterns ubiquitous in nature – emerging in physical, as well as biological phenomena ranging from acoustics, optics, electro-magnetic interactions to morphogenesis.

## Mental states

The relation between brain and mind remains one of the outstanding questions in science (Crick & Koch, 2003). How does our subjective mental experience relate to our neural activity? In this review, we firstly identify different mental states based on their key characteristics in terms of subjective experience and later review the neural correlates of these mental states in terms of temporal, spatial and neurophysiological variations in neural activity.

*Consciousness* although most familiar and intimate to all of us, has been rather ambiguously defined as a scientific subject (Heine et al., 2012; Zeman, 2001). Within the scope of this review, we define consciousness as the mental state that is accompanied by a stream of personal and subjective experiences. Besides during wakefulness, a stream of vivid experiences can also occur in dreaming, meditative states and drug-induced altered states of consciousness. Hence, we consider these states as non-ordinary states of consciousness.

Among various pharmacological substances, a particular class called psychedelics; i.e. psilocybin - the main psychoactive compound in magic mushrooms, lysergic acid diethylamide (LSD), MDMA, ketamine, mescaline and ayahuasca, stand out with their ability to alter consciousness and cognition by introducing an unusual richness to the conscious experience (Carhart-Harris, Erritzoe, et al., 2012a; Carhart-Harris et al., 2014; Kaelen et al., 2015). Due to this enhancement of the conscious experience, the *psychedelic state* is considered to be an “expanded” or “elevated” state of consciousness (Schartner, Carhart-Harris, Barrett, Seth, & Muthukumaraswamy, 2017). A growing body of research is now directed towards understanding the neural markers (Carhart-Harris, Erritzoe, et al., 2012a; Carhart-Harris, Muthukumaraswamy, et al., 2016b; Fernanda Palhano-Fontes, 2015) and mechanisms (Carhart-Harris, 2014) underlying such enhanced and flexible state of consciousness as well as its therapeutic potential for neuropsychiatric disorders (Carhart-Harris, Bolstridge, et al., 2016a).

Unlike waking consciousness and psychedelic states, *sleep state* can be accompanied by both, absence and presence of conscious experience (Siclari et al., 2017). Dream state, where a stream of vivid experiences that can be recalled upon awakening occur, composes the conscious experience in the sleep state, whereas deep sleep in the absence of any dream experience is associated with loss of consciousness. Traditionally, sleep is categorised into two main components, rapid eye movement (REM) sleep and non-REM (NREM) sleep, where NREM sleep can be further divided into 4 stages (N1, N2, N3 and N4) (Rechtschaffen & Kales, 1968). Generally, N1 and N2 are characterized as light sleep, whereas N3 and N4 compose the slow-wave/deep sleep stages.

Loss of consciousness induced by sleep or anaesthesia is a gradual phenomenon (Deco, Hagmann, Hudetz, & Tononi, 2013a) characterized by progressive disengagement from the external world (Deco, Hagmann, Hudetz, & Tononi, 2013a; Larson-Prior et al., 2009) as well as by reduced self-awareness (Samann et al., 2011) followed by a sudden disappearance of both, awareness of the external world as well as the self. A state with the loss of awareness of the self and the environment, induced physiologically (e.g. sleep) or pharamacologically (e.g. anaesthesia) is considered not conscious. Although various other mental states caused by disorders of consciousness such as vegetative and comatose states, psychosis, epileptic seizures can also be identified and individually characterized, in this review we focus on the neural correlates of *wakefulness*; *psychedelic state* (an enhanced state of consciousness) and physiological as well as pharmacological loss of consciousness due to *sleep* and *anaesthesia*.

## Neural correlates of mental states

### Time: Oscillatory correlates of mental states

It has long been known that the amplitude and the frequency of the neural oscillations are altered in different mental states. After the first discovery of low amplitude, “desynchronized” alpha waves (8-12 Hz) in electroencephalography (EEG) recordings of conscious wakeful humans (Berger, 1929), various oscillatory patterns have been identified as the neural correlates of different mental states. In loss of consciousness such as in deep sleep state, although the mean neuronal firing rate is close to those that occur during quiet wakefulness (Massimini, 2005), a transition occurs from the low amplitude, high-frequency alpha oscillations (8-12 Hz) to high-amplitude, low-frequency delta waves (0.5Hz-4Hz) (Simon & Emmons, 1956). Moreover, the two sleep stages, REM and NREM have also substantially different temporal signatures in EEG; NREM is mainly accompanied by slow waves and sleep spindles, whereas REM sleep is characterized by the wakefulness-like, low amplitude, high frequency oscillations. The temporal correlates of dreaming have also been found to closely resemble that of waking conscious state; i.e. globally activated, high-frequency EEG activity. Initially, dreams have been thought to occur only in the rapid-eye-movement (REM) sleep. However, recent studies have discovered that a local decrease in low-frequency EEG activity in the posterior cortex is associated with reports of dream experiences upon awakening during both NREM and REM sleep, whereas a local increase in low frequency activity correlates with the absence of experience (Siclari et al., 2017). These findings indicate that not only the temporal changes, but rather spatially localized changes in temporal activity can characterize presence and absence of conscious experience in the sleep state.

Similarly, neural correlates of anaesthesia-induced loss of consciousness are not only limited to temporal changes in neural oscillations. Recent studies discovered that propofol-induced loss of consciousness was accompanied by the loss of spatially coherent occipital alpha oscillations, and simultaneously by the appearance of spatially coherent frontal alpha oscillations (Purdon et al., 2013). Other EEG studies focusing on evoked cortical responses found that cortical activity patterns after transcranial magnetic stimulation (TMS) during loss of consciousness were spatially more localized compared to the cortical response in a waking conscious state, indicating a reduced integration of cortical activity (Casali et al., 2013) and eventually a breakdown of cortical connectivity in loss of consciousness (Massimini, 2005).

In the psychedelic state, decreased oscillatory power has been observed in the alpha-band (8-13 Hz) in EEG (Eduardo Ekman Schenberg, 2015) after ayahuasca-intake; within low frequency range (0.01-0.1 Hz) in fMRI (Tagliazucchi, Carhart-Harris, Leech, Nutt, & Chialvo, 2014) and within the 8-100 Hz frequency band in MEG after psilocybin administration (Muthukumaraswamy et al., 2013). Psychedelic state induced by another prototypical psychedelic, LSD, has been reported to cause reduced oscillatory power in four different frequency bands delta (1-4Hz), theta (4-8Hz), alpha (8-15Hz) and beta (15-30Hz) in MEG and lead to globally increased functional connectivity (Tagliazucchi et al., 2016, Carhart-Harris, Muthukumaraswamy, et al., 2016b), especially within high-level association cortices (Tagliazucchi et al., 2016) and within spatially distributed cortical areas composing particular resting state networks.

Taken together, these findings converge to suggest that it is not only the frequency and amplitude of neural oscillations that is altered during a change in mental state, but rather the *spatiotemporal* patterns and the *dynamics* of cortical activity.

#### Box. 1. Glossary

**Mental state:** state of conscious awareness; e.g. conscious waking state, psychedelic state, sleep states (REM and NREM consisting of S1, S2, S3 and S4), anaesthesia-induced loss of consciousness.

**Loss of consciousness:** loss of conscious awareness of one’s surrounding and the self. As the sense of self persists in e.g. during REM sleep and dreaming, the loss of consciousness according to this definition occurs in deep sleep and during anaesthesia.

**Neural correlates of a mental state:** spatiotemporal patterns of neural activity accompanying different mental states.

**Temporal correlates of a mental state:** different frequency and amplitude temporal oscillations measured with different functional neuroimaging techniques such as functional magnetic resonance imaging (fMRI), EEG and magnetoencephalography (MEG).

**Network correlates of a mental state:** spatial patterns of correlated (synchronized) activity emerging on the cortex during a particular mental state.

**Structural connectivity:** structural interconnections between brain regions. Development of non-invasive in-vivo imaging techniques such as the diffusion tensor imaging (DTI) allowed for tracing the interconnections between brain regions revealing a macroscale structural mapping termed human connectome.

**Functional connectivity:** spatial patterns of correlated activity among spatially distributed parts of the cortex. Spontaneous oscillations of brain activity in resting state exhibit distinct correlation patterns, referred to as functional connectivity patterns.

**Resting state networks (RSNs):** network of cortical areas exhibiting correlated spontaneous oscillations in the resting state.

**Functional networks:** network of brain areas showing increased activity during a particular cognitive function. These networks have been identified using various task-based paradigms. The major functional networks of the human brain consist of high level cognitive networks such as dorsal attention network, ventral attention network, control network as well as low-level sensory networks such as visual network, auditory network and somato-sensory-motor network.

**Default mode network:** network of brain areas consisting of consists of the medial prefrontal cortex (mPFC), posterior cingulate cortex (PCC)/precuneus (PCu), inferior parietal lobe, lateral temporal cortex and hippocampal formation, which shows increased activity in the absence of any cognitive task and decreased activity during a task performance.

**Brain dynamics:** spatiotemporal evolution of brain activity. Describing brain dynamics involves describing the temporal evolution of spatial patterns of neural activity.

**Neural field models:** equations governing the spatiotemporal patterns of excitatory and inhibitory neural activity in terms of average synaptic or firing rate of populations of neurons. In their generic form, these models consider the neuronal dynamics to play out on a spatially extended cortical sheet, i.e. a neural field. The most commonly used neural field equations are Wilson-Cowan equations (Wilson & Cowan, 1973) expressing the evolution of neural excitatory and inhibitory neural activity as a set of coupled differential equations, which govern the coupling of the two types, excitatory and inhibitory populations.

### Space: Network correlates of mental states

The remarkable discovery that temporal oscillations in spontaneous brain activity exhibit strong correlations within widely distributed cortical regions revealed highly structured correlation patterns in the absence of any task or behaviour (Fox et al., 2005; Fox & Raichle, 2007). Notably, the topography of these correlation patterns, termed the *
**resting state networks (RSNs)**
*, closely resembles the functional networks of the human brain identified by various sensory, motor, and cognitive paradigms (Fox et al., 2005; Fox & Raichle, 2007) (Figure 1).

Initially discovered in spontaneous slow (<0.1 Hz) fluctuations of the blood oxygen level dependent (BOLD) signal measured with fMRI (Fox et al., 2005), these RSNs have been recently also identified using MEG (de Pasquale et al., 2010; 2012; Spadone, de Pasquale, Mantini, & Penna, 2012) and found to correlate with the time courses of electroencephalography (EEG) microstates, i.e. global brain states occurring in discrete epochs of about 100ms. (Britz, Van De Ville, & Michel, 2010; Musso, Brinkmeyer, Mobascher, Warbrick, & Winterer, 2010), suggesting the RSNs’ neuronal origin and relation to temporal oscillations.

Furthermore, even though the correlated activity has been found to transcend levels of consciousness (Fransson, 2005; Fukunaga et al., 2006; Vincent et al., 2007; (Horovitz et al., 2008; Larson-Prior et al., 2009), recent studies reveal a reduction in the strength of correlations in certain networks and substantial changes in the anatomical configuration (coupling of certain cortical areas) of these networks in loss of consciousness (Boveroux et al., 2010; Deco, Hagmann, Hudetz, & Tononi, 2013a; Greicius et al., 2008; Horovitz et al., 2009; Martuzzi, Ramani, Qiu, Rajeevan, & Constable, 2010; Samann et al., 2011; Schrouff et al., 2011; Stamatakis, Adapa, Absalom, & Menon, 2010).

In particular, one network of brain regions, termed *
**default mode network (DMN)**
*, whose activity actually increases in the absence of cognitively demanding tasks (Buckner, Andrews-Hanna, & Schacter, 2008; Fox & Raichle, 2007), has been of particular interest due to its association with conscious, internally mediated cognitive processes, such self-referential behaviour, moral reasoning, recollection, and imagining the future (Buckner et al., 2008; Raichle & Snyder, 2007; Raichle et al., 2001). However, the association of the DMN with the waking state of consciousness has been contrasted by the surprising discovery of DMN’s equivalent in anesthetized monkeys (Vincent et al., 2007).

Vincent et al. discovered that the correlated activity within the DMN, particularly between the posterior cingulate and precuneus cortex (pC-PCC) and lateral temporoparietal components of DMN persisted in anesthetized monkeys (Vincent et al., 2007). A following study reported that although the functional connectivity of the DMN persisted in conscious sedation in humans, connectivity of the posterior cingulate cortex (PCC), that is also a part of the DMN was significantly reduced during sedation (Greicius et al., 2008). Similarly, Stamatakis et al. found that in propofol-induced moderate sedation in humans, the preserved DMN connectivity was accompanied by significant changes in PCC connectivity, especially in the emergence of the inter-areal connections between PCC and the somato-motor cortex, the anterior thalamic nuclei and the reticular activating system (Stamatakis et al., 2010). Martuzzi et al. reported that although the general functional connectivity in DMN did not significantly differ between awake and anesthetised humans, anesthesia modulated the strength of functional connectivity in a network-specific manner; i.e. while connectivity of primary sensory cortices slightly increased, reduced connectivity was observed in high-order association areas (hippocampus and insula) (Martuzzi et al., 2010). Furthermore, propofol-induced decreases in consciousness linearly correlate with decreased connectivity in the DMN and executive control network – which is usually anti-correlated to the DMN (Boveroux et al., 2010). The spatial extent of these anti-correlations greatly diminished during clinical unconsciousness, due to deep sedation (Barttfeld et al., 2015; Boveroux et al., 2010). During propofol-induced loss of consciousness, Schrouff et al. have found reduced integration (estimated by entropy-based hierarchical clustering) within each functional network as well as between networks (Schrouff et al., 2011). Functional connectivity was particularly affected between parietal areas and frontal or temporal regions during deep sedation induced by propofol (Schrouff et al., 2011). These findings indicate that although the DMN was detectable under anaesthesia, loss of consciousness was associated with widespread changes in functional connectivity patterns throughout the thalamo-cortical system, in particular, changes in higher-order fronto-parietal associative network as well as DMN connectivity seem to play a major role in the different states of consciousness.

Interestingly, activity observed during anesthesia and light sedation (Stamatakis et al., 2010) also shows intriguing similarities with slow-wave sleep (Horovitz et al., 2009), suggesting shared neural mechanisms underlying the fading of consciousness due to sedation and NREM sleep (Brown, Purdon, & Van Dort, 2011). Moreover, inline with the findings on anaesthesia-induced connectivity changes, fMRI studies focusing on the connectivity changes in RSNs during sleep also reported persistent functional connectivity in DMN during light sleep (Horovitz et al., 2008). Early fMRI studies comparing light sleep stages to wakefulness (Fukunaga et al., 2006; Larson-Prior et al., 2009) reported no significant changes in the correlation patterns indicating that the RSNs are maintained during light sleep (Larson-Prior et al., 2009). Subsequent studies reported that functional connectivity not only persisted in the DMN, but in 5 RSNs including sensory networks (visual, auditory, and somatomotor) as well as cognitive networks (dorsal attention, default mode, executive control) networks during light (stage 2) sleep (Larson-Prior et al., 2009). However, later studies with more detailed analysis revealed substantial changes in the coupling of cortical areas in different stages of sleep: Using seed-based correlation analysis, Horovitz et al. report a functional decoupling of the frontal cortex from the rest of the DMN components indicated by significant decrease of the strength of correlations in deep sleep (stages 2-4), although the local coherence of spontaneous activity persisted within each individual brain region composing the DMN (Horovitz et al., 2009). These findings were further confirmed by the independent component analysis (ICA) approach of Samann et al. reporting a stepwise decrease and eventual *decoupling of the frontal cortex (medial prefrontal cortex-mPFC) from the DMN with increasing sleep depth* in non-rapid eye movement (NREM) sleep (Samann et al., 2011). Besides the decoupling between the posterior and anterior midline nodes of the DMN, the authors also observe a breakdown of the cortico-cortical functional connectivity between the DMN and its anti-correlated network, which are associated with internal and external awareness during wakefulness. Considering a decomposition of the DMN components into a core and subsystem^
1
^ Koike et. al. find persistent connectivity in DMN core regions in light and deep NREM sleep as well as REM sleep, whereas substantial changes in the functional connectivity in DMN subsystems have been observed between REM and NREM sleep stages (Koike, Kan, Misaki, & Miyauchi, 2011). Using entropy-based hierarchical clustering, Boly et al. find a profound modification of the hierarchical organization of large-scale networks into smaller independent modules in NREM sleep (Melanie Boly et al., 2012). In particular, the authors report an increase in the interactions within each network in comparison to the interactions between different networks as well as in the total brain integration in NREM sleep compared to wakefulness (Melanie Boly et al., 2012).

Taken together, these studies indicate that, although there is not a binary absence/presence relation between the functional connectivity of the RSNs and consciousness, the integrity of the DMN is dynamically modulated by the level of consciousness. Crucially, while earlier studies suggested that the DMN is preserved in early stages of sleep (Horovitz et al., 2008), in-depth analysis revealed substantial changes in the coupling of cortical areas and altered correlation between DMN components in deeper stages of sleep (Horovitz et al., 2009; Melanie Boly et al., 2012; Samann et al., 2011). In particular, a reduced involvement and eventual decoupling of frontal cortex from the DMN was observed with increasingly deeper stages of sleep (Horovitz et al., 2009; Samann et al., 2011).

Interestingly, decreased functional connectivity between the anterior-posterior nodes of the DMN (mPFC and PCC) has also been reported in psilocybin-induced psychedelic state (Carhart-Harris, Erritzoe, et al., 2012a). This reduced within-network functional connectivity observed in the psychedelic state, was found to be accompanied by increased between-RSN functional connectivity (Roseman, 2014). In particular, significantly increased functional connectivity between DMN and its anti-correlated network termed the task positive network (TPN) has been found in psilocybin-induced psychedelic state (Carhart-Harris, Leech, et al., 2012b). Considering DMN’s correlation with internally mediated processes and TPN’s relation to externally focused attention, this increased DMN-TPN connectivity has been hypothesized to underlie the inability to distinguish between one’s internal world and external environment as experienced during psychosis (Carhart-Harris, Leech, et al., 2012b). Another prototypical psychedelic, LSD, has been found to lead to globally increased functional connectivity (Tagliazucchi et al., 2016, Carhart-Harris, Muthukumaraswamy, et al., 2016b), especially within high-level association cortices overlapping with the default-mode, salience, and fronto-parietal attention networks as well as in the thalamus (Tagliazucchi et al., 2016). Particularly, within the DMN, reduced oscillatory power has been observed with MEG (Muthukumaraswamy et al., 2013) and fMRI signals after psilocybin administration (Tagliazucchi et al., 2014); and in the MEG-alpha frequencies (Carhart-Harris, Muthukumaraswamy, et al., 2016b) in the LSD state, where this reduced power has been found to correlate with the experience of ego dissolution, e.g. the dissolution of the subjective self (Carhart-Harris, Muthukumaraswamy, et al., 2016b). Interestingly, a connectome harmonic decomposition, as we discuss here, when applied to fMRI data in the LSD state reveals a frequency-selective expansion and dynamical re-organization of the repertoire of these harmonic brain states at the edge of criticality – transition between order and chaos (Atasoy et al., 2017).

Furthermore, a greater diversity in the repertoire of the functional connectivity patterns in fMRI was observed in the brain under psilocybin together with an increase in the rate at which this repertoire is examined by the brain dynamics (Tagliazucchi et al., 2014). In line with these findings, significant changes in the repertoire of functional connectivity patterns has also been found between wakefulness and anaesthesia-induced loss of consciousness. When analyzed in terms of brain states defined as dominant recurrent patterns of functional connectivity, wakefulness is characterized by the dynamical exploration of a rich, flexible repertoire of brain states compared to anesthesia-induced loss of consciousness (Barttfeld et al., 2015). This opposite trend of enhanced vs. reduced diversity of the functional connectivity patterns observed in psychedelic state vs. anesthesia-induced loss of consciousness points out the significance of the dynamical repertoire of brain states as signatures of different mental states (Atasoy et al., 2017).

These experimental findings are also in agreement with the theoretical propositions stating conscious phenomenology is related to a distributed neural process that is both highly integrated and highly differentiated (Tononi & Edelman, 1998) indicating the need for a large repertoire of highly differentiated brain states. In fact, dynamical systems such as the brain maximize their state repertoire when they approach criticality; i.e. transition between order and chaos, which has also been proposed to be the neural mechanism underlying conscious wakefulness as well as psychedelic experience (Carhart-Harris, 2014). Recently, it has also been pointed out that different mental states may be understood as variations of neural activity along multiple dimensions, instead of just one dimension (variable) (Bayne, Hohwy, & Owen, 2016), hence, rendering the attempt of modelling mental states (or states of consciousness) in one dimensional terms not plausible (Bayne et al., 2016).

Here we introduce a framework for understanding the notion that changes in mental states relate to variations of neural activity along multiple dimensions, instead of just one dimension, which suggests that these various dimensions, may in fact correspond to different harmonic brain modes or different combinations of these harmonic modes. We also show how these harmonic brain modes naturally self-organize from neurophysiological interactions.

### Neurophysiological correlates of mental states

The spatiotemporal patterns of cortical activity at the whole-brain level are thought to emerge from the *local cortical-dynamics* and *cortico-cortical interactions* constrained by the anatomical connectivity of the human brain. The local cortical dynamics involve the interplay of excitation; for instance mediated by glutamatergic principal cells, and inhibition; for instance mediated *γ*-aminobutyric acid GABAergic interneurons (Isaacson & Scanziani, 2011).

The spatial propagation of these interactions between excitatory and inhibitory neural populations locally on the cortical grey matter, as well as between cortical regions connected through the long-distance white matter connectivity of the human brain gives rise to the emergence of spatiotemporal patterns of cortical activity. Thus, functional networks as well as oscillations of neural activity emerge as collective behaviour of individual neurons regulated by neurophysiological variables; i.e. interplay between excitation and inhibition. As such, the same anatomical structure, i.e. human brain connectivity gives rise to a variety of oscillatory patterns and networks simply by alteration of excitatory and inhibitory activity.

Recently, development of novel structural neuroimaging technologies, such as diffusion tensor imaging, have enabled the tracking of the long-distance white matter fibres of the human brain providing a comprehensive structural description of brain’s network architecture, termed *
**human connectome**
* (Sporns, Tononi, & Kötter, 2005). Incorporating the human connectome into theoretical and computational models of neural dynamics has provided valuable insights into the link between anatomical structure of the human brain and RNS dynamics (Atasoy et al., 2016; Deco et al., 2009b; Deco, Jirsa, & McIntosh, 2013b; Honey et al., 2009). Furthermore, such whole-brain computational models also enable the investigation of cortical dynamics accompanying different mental states.

While the exact neural mechanisms underlying the loss and recovery of consciousness remain largely unknown, converging neurophysiological evidence suggests that transition to an unconscious state is accompanied by increasing inhibitory and/or decreasing excitatory activity (Brown, Lydic, & Schiff, 2010; Tononi & Koch, 2008). In contrast to loss of consciousness, the psychedelic state, an enhanced state of consciousness, is postulated to result from increased cortical excitation via the stimulation of 5-HT2A receptors caused by psychedelic drugs (Glennon, Titeler, & McKenney, 1984). All psychedelic drugs are agonists at the serotonin 2A receptor (5-HT2AR) leading to increased 5-HT2AR signaling (Glennon et al., 1984). In the cortex, this receptor is most densely expressed postsynaptically in layer 5 pyramidal neurons, which are large excitatory neurons. Hence increased 5-HT2AR stimulation by the psychedelic substance results in increased cortical excitation (Celada, Puig, & Artigas, 2013). Although different anesthetics, psychedelics and sleep may affect neurophysiology through different neuronal pathways and mechanisms, converging evidence suggests that *drug- or sleep-induced loss of consciousness is associated with increasing inhibitory or decreasing excitatory activity, whereas psychedelic-induced enhanced state of consciousness results from increasing excitatory activity*. Hence, the contrast of these mental states; i.e. loss vs. enhancement of conscious experience, is also reflected in the activation of opposite neurophysiological mechanisms; i.e. decreasing vs. increasing cortical excitation.

Here we demonstrate how the above reviewed, seemingly unrelated findings on the neural correlates of consciousness; changes in neural oscillations, RSN configuration, the dynamical repertoire of brain states and in neurophysiology accompanying different mental states (wakefulness, sleep, anaesthetised and psychedelic states) *can be unified* and understood within one framework expressing the neural activity *as combinations of harmonic brain modes*. To this end, we first introduce the concept of elementary brain modes as the building blocks of spatiotemporal neural activity. Crucially, the particular definition of the elementary brain modes as connectome harmonic, proposed here, links both, the functional connectivity patterns as well as the cortical oscillations to the particular structural connectivity of the human brain, hence revealing the relation between spatial and temporal aspects of cortical activity patterns and human brain anatomy. Within the introduced framework, we then illustrate how a simple change in the excitation/inhibition (E/I) balance results in the change of temporal as well as network-correlates of mental states simply by activating and deactivating a different combination of these elementary brain modes.

## Linking space and time of brain dynamics via harmonic brain modes

The RSNs, i.e. networks of distributed brain regions with synchronized spontaneous oscillations, are revealed by estimating patterns of functional connectivity, e.g. by estimating the correlation value between the temporal activity of a seed-location and that of the rest of the cortex (Fox & Raichle, 2007). Hence, these spatial correlation patterns are intrinsically linked to spontaneous temporal oscillations of neural activity. However, as these spontaneous oscillations contain various temporal frequencies, the resulting spatial patterns of functional connectivity also consists of correlation patterns corresponding to multiple temporal frequencies.

Here, we introduce the idea that just like the different combinations of chemical basis in DNA comprise the fundamental elements of different genes or elementary particles form the building blocks of various atoms and molecules, complex spatiotemporal patterns of neural activity can also be decomposed into simple, elementary building blocks, i.e. *
**elementary brain modes**
*. Taking into account the intrinsic relation between the temporal oscillations and the functional connectivity patterns they compose, an efficient definition of elementary brain modes would be patterns of fully synchronous neural activity emerging for different temporal frequencies. In fact, recently it has been shown that for each temporal frequency, the accompanying fully synchronous cortical pattern can be estimated from the structural connectivity of the human brain via the mathematical framework of harmonic modes - a fundamental principle governing a multitude of physical and biological phenomena (Atasoy et al., 2016).

Harmonic patterns are ubiquitous in nature emerging as building blocks of pattern formation in various physical and biological phenomena: standing wave patterns emerging in sound-induced vibrations of a guitar string or a metallic plate (first demonstrated as complex sand patterns by Chladni (Chladni, 1830)), patterns of ion motion emerging from electro-magnetic interactions (Britton et al., 2012; Roos, 2012), electron wave function of a free particle given by time-independent Schrödinger equation (Moon et al., 2008; Schrodinger, 1926) and even in patterns emerging in complex dynamical systems such as the reaction-diffusion models (Murray, 1990; Xu, Vest, & Murray, 1983). Mathematically, these harmonic patterns are given by the eigenfunctions of the Laplace operator Δ, which lies at the heart of theories of heat, light, sound, electricity, magnetism, gravitation and fluid mechanics (Stewart, 1999):

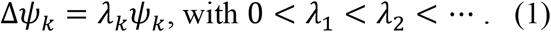

Here we draw on the standing waves emerging in resonance phenomena to gain insights into the harmonic patterns and temporal oscillations. In principle, every physical system has a spectrum of preferred frequencies, called *natural frequencies*, at which the system tends to oscillate. When a physical system such as Chladni’s metal plate or a musical instrument is excited with one of its natural frequencies, the system resonates and a sudden change in state occurs leading the system to a dynamic stable state, called the *eigenmode* or the natural mode of vibration. In each eigenmode, a characteristic standing wave emerges in the system and is maintained by the oscillations in the corresponding natural frequency. In other words, the standing wave is formed and sustained by the oscillation of each spatial location with the same natural frequency, which is synchronized throughout the complete system; e.g. metal plate. This intrinsic relation between the standing wave patterns and the corresponding natural frequency of the temporal oscillations is also reflected in the mathematical description of this phenomenon; i.e. the eigenfunctions of the Laplace operator Δ, i.e. the harmonic patterns *ψ*
_k_, yield the shape of the standing wave pattern adapted to the particular geometry of the underlying domain, and the corresponding eigenvalues *λ*
_k_ relate to the activated natural frequency. Hence, the eigendecomposition of the Laplace operator, Eq. (1) also known as the Helmholtz equation, links the temporal oscillations, the spatial patterns of synchrony and the particular geometry where the standing waves emerge.

Remarkably, when applied to a one-dimensional domain with cyclic boundary conditions; i.e. to a ring, these harmonic patterns constitute the well-known Fourier basis, whereas on a sphere they yield the spherical harmonics. Notably, recent work has shown that the extension of these harmonic patterns to the particular structural connectivity of the human brain, the human connectome, predicts the resting state networks (Atasoy et al., 2016). This notable finding suggests that the same fundamental principles governing harmonic patterns in other natural phenomena also underlie the cortical patterns emerging from the collective dynamics of neural activity (Atasoy et al., 2016).

In this review, we emphasize another key characteristic of connectome harmonics: Due to their orthogonality, the set of all connectome harmonics provides a new function basis, which is by definition, an extension of the well-known Fourier basis to the structural connectivity of the human brain. The same way that any continuous signal can be reconstructed from the Fourier basis, any pattern of cortical activity can also be decomposed and reconstructed from the set of connectome harmonics. Thus, the set of connectome harmonics provides *a new frequency-specific language to describe cortical activity*, where each connectome harmonic pattern corresponds to a frequency-specific building block – an elementary brain mode - composing complex patterns of cortical activity. Moreover, due to their implicit relation to resonance phenomena, where standing wave patterns are intrinsically linked to natural frequencies as well as to the geometry of the underlying domain, the language of connectome harmonics also establishes a natural link between the functional connectivity patterns, the temporal oscillations and the anatomy of the human brain.

**Figure 2.**
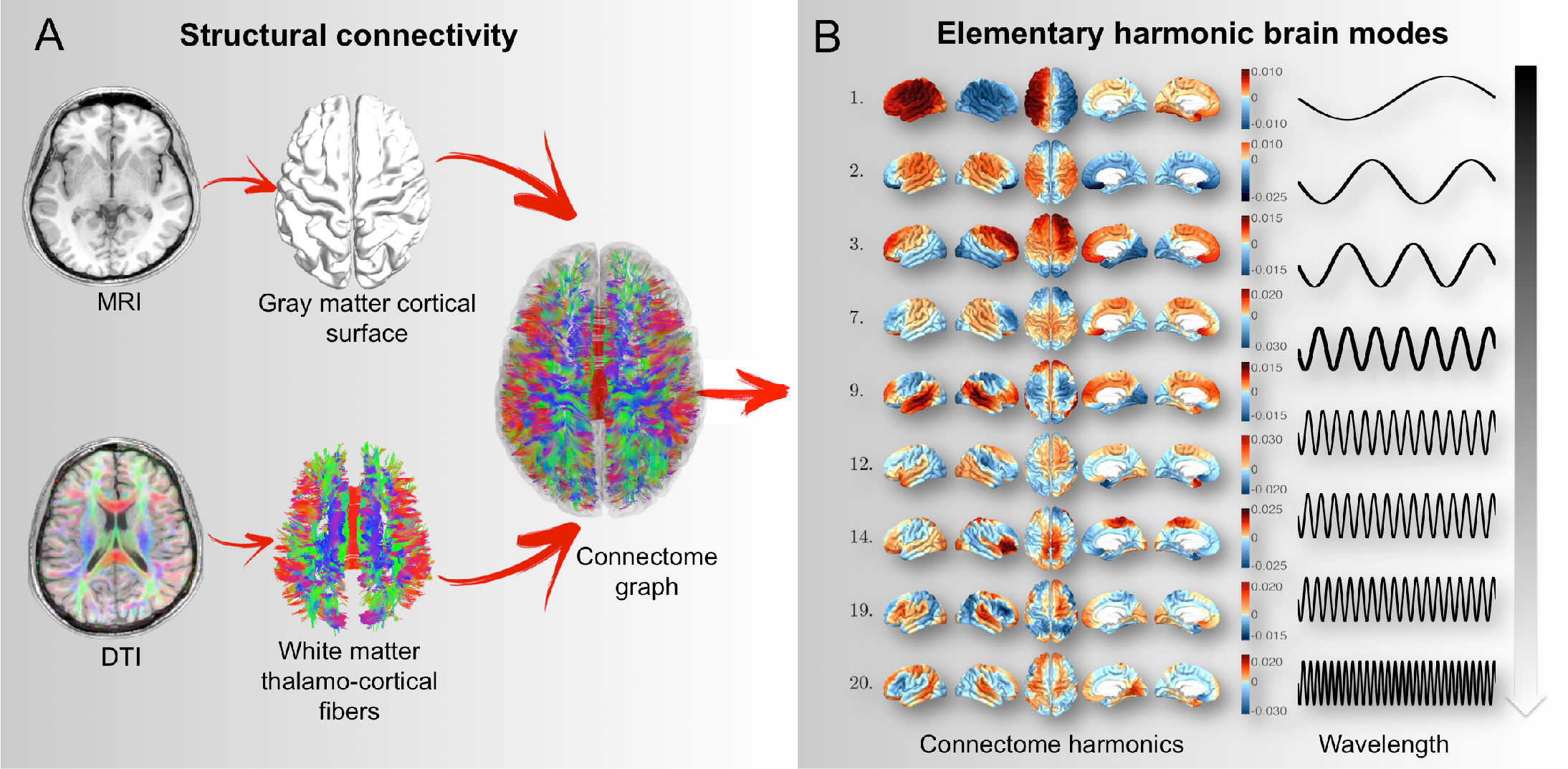
**A**) Structural connectivity of the human brain defined as the combination of local cortical, gray matter connections. **B**) Elementary harmonic brain modes defined as fully synchronous patterns of neural activity are estimated as the harmonic modes of structural connectivity; i.e. connectome harmonics.

Remarkably, when expressed in this new harmonic language, different RSNs have been found to match different parts of the connectome harmonic spectrum (Atasoy et al., 2016). Furthermore, connectome harmonics have been shown to naturally self-organize from the interplay between neural excitation and inhibition (Atasoy et al., 2016). Next we demonstrate how seemingly unrelated experimental findings on the neural correlates of different mental states can be unified and understood in terms of activation and deactivation of these harmonic brain modes – connectome harmonics – and how this activation and deactivation is mediated crucially by the excitation-inhibition balance.

**Figure 3.**
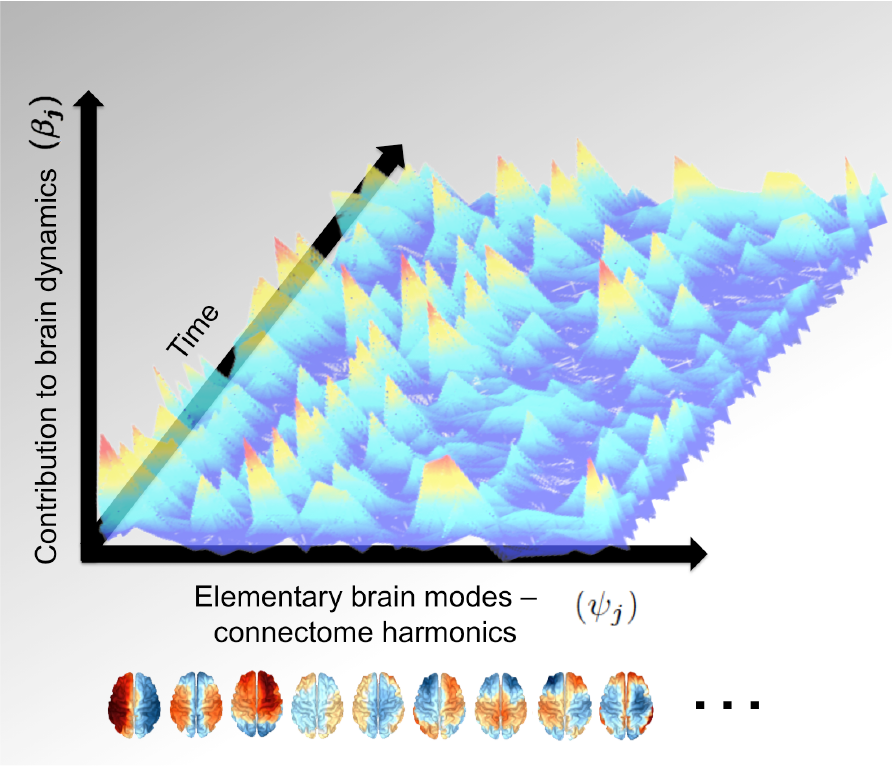
Projection of spatiotemporal patterns of neural activity reveals the contribution (*β*
_
*j*
_) of each connectome harmonic (*ψ*
_
*j*
_) - elementary harmonic brain mode - to brain dynamics.

## A unifying framework for neural correlates of mental states

In the previous sections, we reviewed three different aspects of the neural correlates of mental states: 1) neural oscillations 2) functional connectivity patterns, and 3) the underlying neurophysiological interactions. In the last Section we demonstrated how the temporal oscillations and spatial functional connectivity patterns are linked to each other, as well as to neuroanatomy through the principle of harmonic waves; i.e. via connectome harmonics. Now, we utilize an extension of neural field models, a well-studied mathematical model to describe neural activity via the interactions between excitation and inhibition, to demonstrate the link between the self-organization of connectome harmonics and neurophysiological changes accompanying various mental states.

Neural field modes are a set of coupled differential equations, which govern the coupling of the two types, excitatory and inhibitory neuronal populations in terms of average synaptic or firing rate. In their generic form, these models assume that neuronal dynamics to play out on a spatially extended cortical sheet, i.e. a neural field. The most commonly used neural field equations, Wilson-Cowan equations (Wilson & Cowan, 1973), have been successfully applied to describe the evolution of neural excitatory and inhibitory neural activity in the thalamo-cortical system (Wilson & Cowan, 1973), formation of ocular dominance patterns in the visual cortex (Swindale, 1980) and patterns of visual hallucinations (G. B. Ermentrout & Cowan, 1979; Rule, Stoffregen, & Ermentrout, 2011). Initially introduced to describe neural activity at the mesoscopic scale, extension of neural field models to the full structural connectivity of the human connectome has been recently shown to account for the emergence of spontaneous oscillations and functional connectivity patterns in the macroscopic scale (Atasoy et al., 2016). Here, we utilize this connectome-wide neural field model to investigate which connectome harmonics emerge for a particular excitation/inhibition balance while varying the corresponding parameters in the neural field model according to neurophysiological changes in different mental states.

In neural field models, neural activity is described in terms of activity of two types of populations; activity of excitatory neurons; e.g. mediated by glutamatergic principal cells, exciting excitatory (EE) as well as inhibitory neural populations (EI) and inhibition; e.g. mediated by *γ*-aminobutyric acid GABAergic interneurons, inhibiting excitatory (IE) as well as inhibitory neuronal populations (Figure 4A,B). While in their generic form these two types of activities propagate only locally simulated by the application of a diffusion kernel, in their connectome-wide extension, this spatial propagation occurs not only locally but also through the long-distance white-matter fibers, following a diffusion process on the human connectome (Figure 4A,B). Due to the orthogonality of connectome harmonics and the intrinsic relation between harmonic functions and diffusion equation, the connectome-wide neural field equations can be reformulated in terms of their connectome harmonic expansion (Figure 4C), (Atasoy et al., 2016). This decomposition of the neural field equations into connectome harmonics reveals the contribution of individual connectome harmonics over time to the spatiotemporal dynamics of brain activity. In other words, the neural field equations compose complex oscillatory patterns of cortical activity by letting certain connectome harmonic patterns grow (“activate”) and forcing others to die out (“deactivate”), where the choice of which connectome harmonics are activated and deactivated depends on the excitatory and inhibitory parameters of the model.

**Figure 4.**
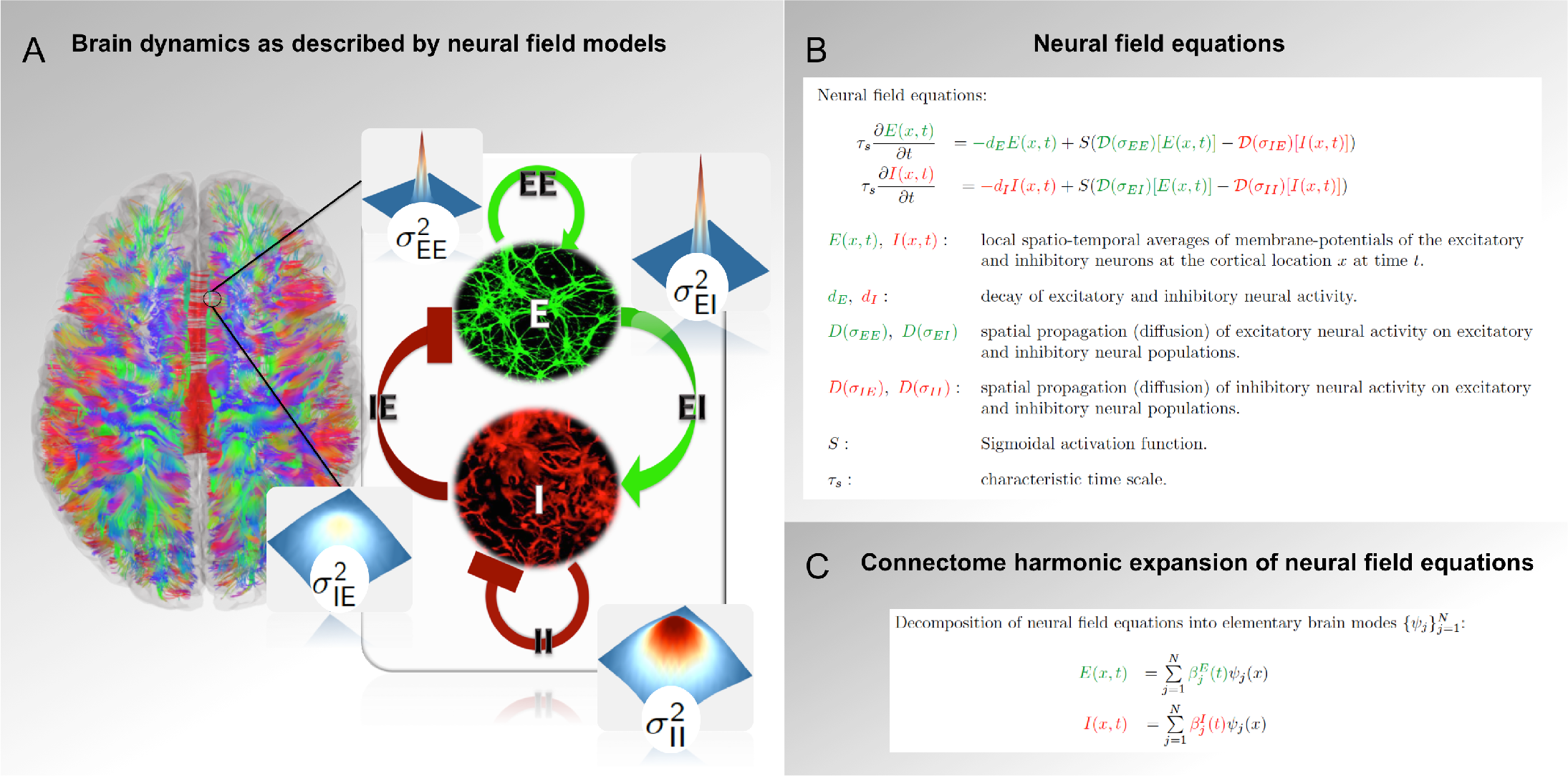
Illustration of the self-organization of the harmonic brain modes from the interplay between neural excitation and inhibition and the decomposition of spatiotemporal brain dynamics into harmonic brain modes. (**A**) Extension of neural field models to the full structural connectivity of human connectome. (**B**) Wilson-Cowan equations. (**C**) Decomposition of Wilson-Cowan equations into connectome harmonics.

Evaluation of the neural field parameters governing the spatial propagation of excitatory EE, EI and inhibitory activity IE and II (Figure 5A) reveals that a decrease in excitation (decrease of σ_EE_ and σ_II_) and increase in inhibition (σ_EI_ and σ_IE_) lead to a *collapse of the repertoire of connectome harmonics to a narrow range of low frequencies* (Figure 5B). Furthermore, this change in the repertoire of active connectome harmonics is also accompanied by a *slowing down of the temporal oscillations* and *decoupling of the frontal cortex from the DMN* (Figure 5C). Remarkably, these findings exactly reflect the neural correlates of sleep- and anesthetic-induced loss of consciousness: In terms of neurophysiological changes, although at the cellular level various anaesthetics and sleep may act differently, drug- or sleep-induced loss of consciousness is associated with increasing inhibitory or decreasing excitatory activity (Tononi & Koch, 2008). On a network level, substantial changes occur in the anatomical configuration of the RSNs and the functional connectivity between the anterior-posterior parts of the DMN breaks down (Table 1). Moreover, a collapse in the repertoire of functional connectivity patterns is observed in anesthesia (Barttfeld et al., 2015). Finally, these neurophysiological and network level changes are also accompanied by a transition from the low amplitude, high-frequency patterns to low-frequency coherent oscillations in cortical activity in loss of consciousness (Casali et al., 2013; Tononi & Koch, 2008); e.g. as observed during the emergence of delta waves in deep sleep.

**Figure 5.**
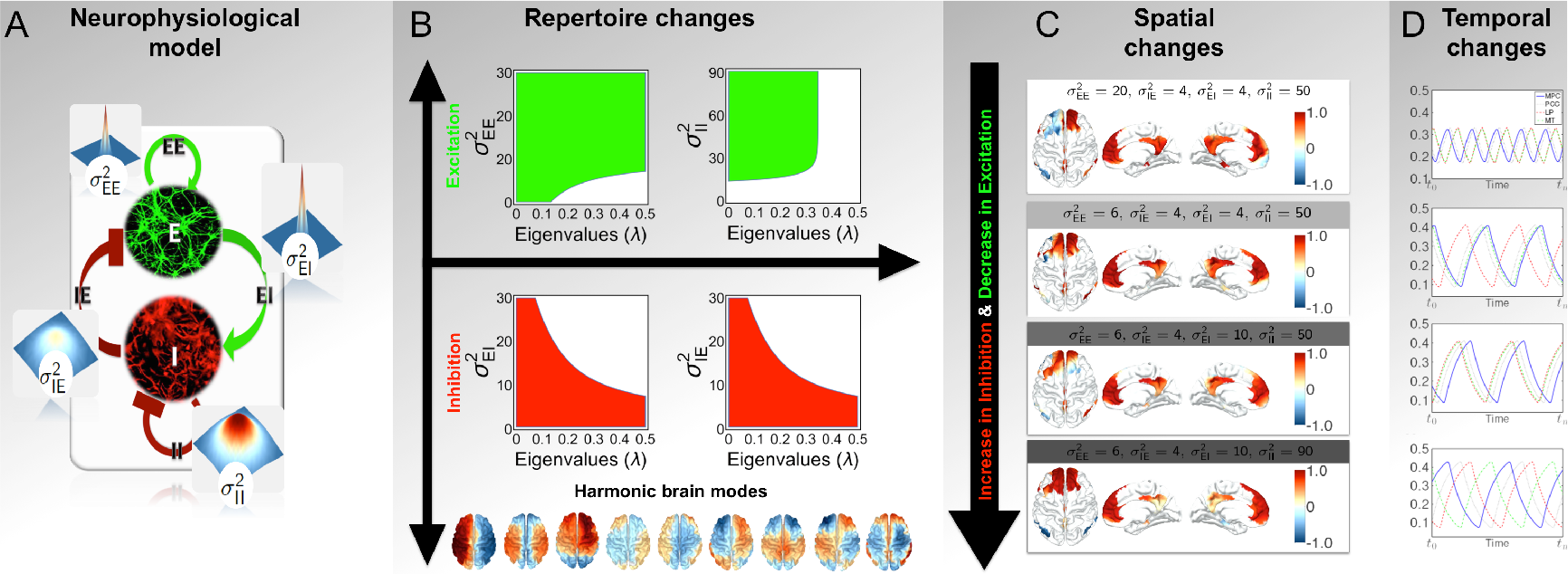
Unifying framework for neural correlates of loss and recovery of consciousness. (**A**) Neural field model of neurophysiological interactions between excitatory and inhibitory neuronal populations. **B**) For increased inhibition and decreased excitation, the repertoire of active connectome harmonics becomes limited to narrow range of low frequencies, as also seen in loss of consciousness. Opposite effect, extended repertoire of connectome harmonics is seen for increased excitation, as also observed in the psychedelic state. Increasing inhibition and decreasing excitation in the neural field model results in (**C**) decoupling of the frontal cortex from the rest of the DMN (correlations are shown for a seed location within the DMN) and (**D**) slowing down of the cortical oscillations as observed during loss of consciousness.

**Table 1:** NETWORK LEVEL CHANGES IN LOSS OR ENHANCEMENT OF CONSCIOUSNESS. The colors of the rows indicate different mental states. Sleep, anesthesia and psychedelic states are marked as blue, green and o ange, respectively. The darkness of the blue and green indicate the depth of sleep and the depth of anesthesia respectively.

| Study | State of consciousness | Imaging Modality | Neural correlates |
| --- | --- | --- | --- |
| (Fukunaga et al., 2006) | Light sleep | fMRI | Persistent patterns of correlated fluctuations. |
| (Horovitz et al., 2008) | Light sleep | fMRI | Persistent correlated fluctuations in DMN. |
| (Larson-Prior et al., 2009) | Light sleep | fMRI | Persistent functional connectivity in 5 RSNs including sensory networks (visual, auditory, and somatomotor) as well as cognitive networks (dorsal attention, default mode, executive control) networks during light (stage 2) sleep. |
| (Horovitz et al., 2009) | Deep sleep<br>Stages 2/3/4 | fMRI | Decoupling of the brain's default mode network during deep sleep. |
| (Samann et al., 2011) | NREM sleep<br>Stage 1,2, slow wave sleep | fMRI | Breakdown of cortico-cortical functional connectivity, particularly between the posterior and anterior midline node of the DMN and the DMN and the ACN (Samann et al., 2011). |
| (Koike et al., 2011) | Light NREM (Stage 2)<br>Deep NREM(Stage 3/4)<br>REM sleep | fMRI | Substantial changes in the functional connectivity in DMN subsystem detected after decomposing the DMN components into a core and subsystem: persistent connectivity in DMN core regions in light and deep NREM sleep as well as REM sleep. |
| (Melanie Boly et al., 2012) | Light NREM (Stage 2)<br>Deep NREM (Stage 3/4) | fMRI | Modification of the hierarchical organization of large-scale networks into smaller independent modules during NREM sleep, was found using information theoretic measures (entropy) and functional clustering index. |
| (Vincent et al., 2007) | Anesthesia (monkeys) | fMRI | Persistence of correlated activity within the DMN, particularly between the posterior cingulate/precuneus cortex (pC/PCC) and lateral temporoparietal components of DMN observed in anesthetized monkeys. |
| (Greicius et al., 2008) | Light, conscious sedation | fMRI | Persistent DMN connectivity detected during conscious, light sedation.<br>Significantly reduced functional connectivity found in the posterior cingulate cortex during conscious sedation. |
| (Stamatakis et al., 2010) | Low and moderate sedation | fMRI | Persistent DMN connectivity detected during low and moderate sedation.<br>Significantly increased functional connectivity found between the posterior cingulate cortex and the motor/somatosensory cortices, the anterior thalamic nuclei, and the reticular activating system, which are not part of the DMN. |
| (Boveroux et al., 2010) | Deep sedation, clinical unconsciousness | fMRI | Propofol-induced decrease of consciousness linearly correlates with decreased connectivity in the DMN and executive control network – which is usually anti-correlated to the DMN. The spatial extent of these anti-correlations greatly diminished during clinical unconsciousness due to deep sedation. |
| (Martuzzi et al., 2010) | Deep sedation, clinical unconsciousness | fMRI | Alteration of the strength of functional connectivity during anesthesia in a network-specific manner. |
| (Schrouff et al., 2011) | Propofol induced<br>Deep sedation, clinical unconsciousness | fMRI | Reduced integration (estimated by entropy-based hierarchical clustering) within each functional network as well as between networks during propofol-induced loss of consciousness. Functional connectivity was particularly affected between parietal areas and frontal or temporal regions during deep sedation induced by propofol. |
| (Carhart-Harris, Erritzoe, et al., 2012a) | Psilocybin-induced psychedelic state | fMRI | Decreased functional connectivity between anterior-posterior nodes of the DMN (mPFC and PCC).<br>Decreases activity in ACC and mPFC. |
| (Carhart-Harris, Leech, et al., 2012b) | Psilocybin-induced psychedelic state | fMRI | Increased functional connectivity between DMN and its anti-correlated task positive network (TPN). |
| (Muthukumaraswamy et al., 2013) | Psilocybin-induced psychedelic state | MEG | Decreased oscillatory power within the DMN measured with MEG.<br>Reduced spontaneous oscillatory power in posterior association cortices within 1-50 Hz and in frontal association cortices within the 8-100 Hz frequency band. |
| (Roseman, 2014) | Psilocybin-and MDMA-induced psychedelic state | fMRI | Increased between-RSN functional connectivity under psilocybin.<br>MDMA had a notably less marked effect on between-RSN functional connectivity. |
| (Tagliazucchi et al., 2014) | Psilocybin-induced psychedelic state | fMRI | Decreased oscillatory power in low frequency range (0.01-0.1 Hz) measured with fMRI within DMN, control and attention networks.<br>Increased diversity in the repertoire of functional connectivity patterns in fMRI and increased exploration rate of this repertoire under psilocybin. |
| (Fernanda Palhano-Fontes, 2015) | Ayahuasca-induced psychedelic state | fMRI | Decreased functional connectivity between PCC and precuneus, parts of the DMN.<br>Increased activity in DMN including Posterior Cingulate Cortex (PCC)/Precuneus and the medial Prefrontal Cortex (mPFC).<br>No significant change in the anti-correlations between DMN and task positive network. |
| (Eduardo Ekman Schenberg, 2015) | Ayahuasca-induced psychedelic state | EEG | Decreased oscillatory power in the alpha-band (8-13 Hz). |
| (Tagliazucchi et al., 2016) | LSD-induced psychedelic state | fMRI | Globally increased functional connectivity throughout the cortex, in particular within high-level association cortices (partially overlapping with the default-mode, salience, and fronto-parietal attention networks) and the thalamus. |
| (Carhart-Harris, Muthukumaraswamy, et al., 2016b) | LSD-induced psychedelic state | fMRI, MEG | Decreased functional connectivity in DMN, as well as various other RSNs (medial visual, lateral visual, occipital pole, auditory, sensorimotor, parietal cortex, dorsal attention, left frontoparietal and right frontoparietal networks).<br>Decreased oscillatory power in four frequency bands delta (1-4Hz), theta (4-8Hz), alpha (8-15Hz) and beta (15-30Hz). In particular decreased oscillatory alpha power in DMN correlates with the experience of ego-dissolution.<br>Decreased functional connectivity and decreased oscillatory alpha power in DMN, as well as decreased functional connectivity between parahippocampus and retrosplenial cortex correlated with the experience of ego-dissolution. |
| (Atasoy et al., 2017) | LSD-induced psychedelic state | fMRI | Frequency selective increase in the repertoire of harmonic brain states. Increased temporal correlation between the co-activation of different frequency brain states suggesting that the increased repertoire causes a re-organization of brain states rather than a random activation.<br>Enhanced signatures of criticality in terms of power laws suggesting that the re-organization of the active harmonic brain states occurs at the edge of criticality. |

To the contrary, psychedelic state – an enhanced state of consciousness – is neurophysiologically associated with increased excitatory activity mediated by 5-HT2AR signaling (Glennon et al., 1984). Notably, increasing the excitatory parameters in the neural field model enables the activation of a broad range of the connectome harmonic spectrum (Figure 5B). Activation of a greater repertoire of connectome harmonics would also lead to higher variability of the temporal fluctuations due to simultaneous contribution of a larger repertoire of temporal frequencies. Indeed, higher temporal variability of the fMRI BOLD (Tagliazucchi et al., 2014) and the MEG signal (Schartner et al., 2017) have been found in the psychedelic state.

Taken together, these findings demonstrate that cortical dynamics modelled by the connectome-wide neural field models remarkably fit the neurophysiological, network-level and temporal correlates of loss and recovery of consciousness while revealing how these different aspects in fact can be simply linked by the activation and deactivation of connectome harmonics as elementary brain modes. Hence, connectome harmonics introduced here as elementary harmonic brain modes not only unifies seemingly unrelated neural correlates of various mental states, but also suggests a fundamental principle ubiquitously observed in various different natural phenomena as their underlying mechanism.

## Conclusions

Here we reviewed existing findings on the neural correlates of different mental states including wakefulness, sleep- and anaesthesia-induced loss of consciousness and psychedelic states. We characterized the neurophysiological, temporal and network-level correlates of these mental states. We introduced the concept of *elementary harmonic brain modes*; i.e. building blocks of spatiotemporal patterns of neural activity and proposed their estimation as connectome harmonics; i.e. harmonic modes of structural connectivity (Atasoy et al., 2016). We then demonstrated the intrinsic link between connectome harmonics and temporal oscillations and discussed how they naturally self-organize from the interplay between neural excitation and inhibition. These intrinsic links of connectome harmonics to neurophysiology and temporal oscillations renders them uniquely suitable to be defined as the elementary brain modes composing the complex spatiotemporal patterns of brain activity. Just like the musical notes composing a complex musical piece, these harmonic brain modes defined as connectome harmonics yield the frequency-specific building blocks of cortical activity.

Remarkably, when neural dynamics - expressed by the connectome-wide neural field model - are decomposed into different connectome harmonics, the critical relation between excitation-inhibition balance and temporal oscillations fits the neurophysiological changes observed during loss and recovery of consciousness. In other words, during loss of consciousness neural activity becomes locked to a narrow frequency range, whereas in wakefulness a broad frequency range of the connectome harmonic spectrum constitutes the neural activity. A broader range of connectome harmonics are enabled for activation for increased excitation which is known to occur in the psychedelic state (Glennon et al., 1984). Hence, following the musical analogy, consciousness can be compared to a rich symphony played by an orchestra, whereas loss of consciousness would correspond to a limited repertoire of musical note played repetitively.

The framework of elementary harmonic brain modes offers a unifying perspective and explanatory framework revealing the link between various seemingly unrelated findings on neural correlates of consciousness. The proposed framework links the spatial patterns of correlated neural activity, not only to temporal oscillations characteristic of mammalian brain activity but also to brain anatomy and neurophysiology. Hence, this framework goes beyond enabling a new dimension of tools for decomposing complex patterns of neural activity into their elementary building blocks, by also providing a fundamental principle linking space and time in neural dynamics through harmonic waves – a phenomenon ubiquitous in nature.

## Footnotes

1 DMN subsystem is considered part of the DMN, but connectivity within the DMN subsystem is not robust as the DMN core, in other words these components are strongly linked to the core hubs of the DMN but not to each other (Buckner et al., 2008).

